# Reduced spatial integration in the ventral visual cortex underlies face recognition deficits in developmental prosopagnosia

**DOI:** 10.1101/051102

**Authors:** Nathan Witthoft, Sonia Poltoratski, Mai Nguyen, Golijeh Golarai, Alina Liberman, Karen F. LaRocque, Mary E. Smith, Kalanit Grill-Spector

## Abstract

Developmental prosopagnosia (DP) is characterized by deficits in face recognition without gross brain abnormalities. However, the neural basis of DP is not well understood. We measured population receptive fields (pRFs) in ventral visual cortex of DPs and typical adults to assess the contribution of spatial integration to face processing. While DPs showed typical retinotopic organization of ventral visual cortex and normal pRF sizes in early visual areas, we found significantly reduced pRF sizes in face-selective regions and in intermediate areas hV4 and VO1. Across both typicals and DPs, face recognition ability correlated positively with pRF size in both face-selective regions and VO1, whereby participants with larger pRFs perform better. However, face recognition ability is correlated with both pRF size and ROI volume only in face-selective regions. These findings suggest that smaller pRF sizes in DP may reflect a deficit in spatial integration affecting holistic processing required for face recognition.

---

Individuals with developmental prosopagnosia (DP) show impaired recognition of faces with otherwise normal cognitive abilities(1)(2)(3)(4), although in some cases other perceptual deficits can be found(5). DPs have difficulty recognizing familiar faces, struggle to learn new faces, and in extreme cases can fail to recognize members of their own family(3). Although recently discovered(6), and initially considered rare(4), DP appears to be a relatively common disorder with an estimated prevalence of 2-2.5% of the population(7).

DP does not arise from head injury or gross anatomical abnormalities easily observed using magnetic resonance imaging (MRI). Therefore researchers have tested for functional abnormalities in face-selective regions of ventral visual cortex using neuroimaging methods such as functional MRI (fMRI). These core face-selective regions include the occipital face area (OFA/IOG-faces(8)) and the pair of face patches on the fusiform gyrus (pFus and mFus)(9), which are often collectively referred to as the fusiform face area (FFA)(10). A substantial body of research from neuropsychology(11), fMRI(12), and electrical brain stimulation studies(13), indicates these regions are involved in the perception and recognition of faces. Surprisingly, the field has not reached a consensus regarding the existence of differences in core face-selective areas in DPs. Furl et al. found smaller volume of right face-selective fusiform regions in DPs compared to controls, which correlated with face identification ability(14). However, other fMRI studies of DP have failed to find differences in the volume, selectivity, or number of core face-selective regions(15)(16)(17)(18).

Several explanations may account for the lack of consistent differences in the core face-selective areas in DP, none of which are mutually exclusive. Some issues may be methodological, as studies differ in fMRI analysis methods (e.g. spatial smoothing) and sample sizes are often small. Given the complexity of face recognition and the distributed brain network that supports it, there may also be heterogeneity in the causes of DP(4). Some researchers have proposed that DP is associated with differences beyond the core face-selective regions, with recent results linking DP to functional abnormalities in face-selective regions of anterior temporal cortex(18), alterations in white matter adjacent to core face-selective regions(19), or reductions in the structural integrity of large fasciculi connecting the core and extended face network(20).

It is also possible that face selectivity alone does not fully capture the computations that underlie face processing. Normal face recognition is especially dependent on configural/holistic processing(21), which integrates spatially disparate facial features into a complex representation that includes the relative arrangement and spacing of facial features’(22). Behavioral evidence(23)(24)(25)(26)(27), and neuroimaging data(17), shows that many (but not all(28)) DPs have a deficit in holistic processing. Moreover, researchers (ourselves included) have noted that DPs report compensatory strategies relying on local features of a face, such as counting eyebrow hairs(1) when remembering or comparing faces.

We therefore posit that neural computational units (neuron or population of neurons) that process faces holistically, must at minimum access the spatially disparate features of a face. An impairment in face recognition could therefore arise if computational units process only local regions of the visual field. This hypothesis predicts that DPs would show face selectivity due to higher responses to facial features than other stimuli, but neuronal populations in core face-selective regions in DP would process a smaller extent of the visual field compared to typicals(17)(29). Consequently, participants with neural populations that process larger portions of the visual field would perform better in face recognition than those with smaller ones.

We can measure spatial processing in the human brain using population receptive field (pRF) modeling(30)(31). PRFs delineate the portion of the visual field that modulates responses in a voxel. We model the pRF of each voxel as a 2D Gaussian, with parameters defining the position of its center and its width in degrees of visual angle. Although pRF modeling has been primarily used to characterize properties of visual field maps, core face-selective regions also contain neural populations that respond preferentially to particular locations in the visual field(32), and can be modeled using pRFs(33). We hypothesized that the face processing deficits seen in DP are due to smaller pRFs in core face-selective regions compared to typicals(29), which do not allow the typical computations needed for face recognition because of the limited portion of the face being processed.

To test this hypothesis, we used fMRI to define face-selective regions and visual field maps, and estimated pRFs in 7 DPs (Supplemental Figures 1,2) and 15 typical adults. We asked: (1) Do DPs show a deficit in spatial processing as evidenced by smaller pRFs? (2) If DPs have smaller pRFs, are they confined to face-selective regions or are they distributed more widely throughout ventral visual cortex? A recent study based on similar reasoning reported that autistic subjects show larger pRFs in V1-V3(29). (3) Is there a relationship between pRF size and face recognition performance?

## RESULTS

DPs and typicals had broadly similar functional organization in ventral visual cortex(15) (**Figure 1**). DPs had normal eccentricity and polar angle maps and did not differ from typicals in the anatomical location of visual field maps. We defined V1, V2, V3, hV4, and VO1(34)(35), in every hemisphere of every participant in both groups. We also defined three face-selective regions of interest (ROIs) in each hemisphere, one on the inferior occipital gyrus (IOG-faces/OFA), lateral to hV4, and two on the lateral bank of the fusiform gyrus, one in the posterior fusiform (pFus-faces) and one more anterior, neighboring the mid fusiform sulcus (mFus-faces)(9). While not all participants had all face-selective ROIs in both hemispheres, the two groups did not differ in the number of subjects with any given face ROI (Exact p =1 in all cases, Supplementary Table 5).

**FIGURE 1:**
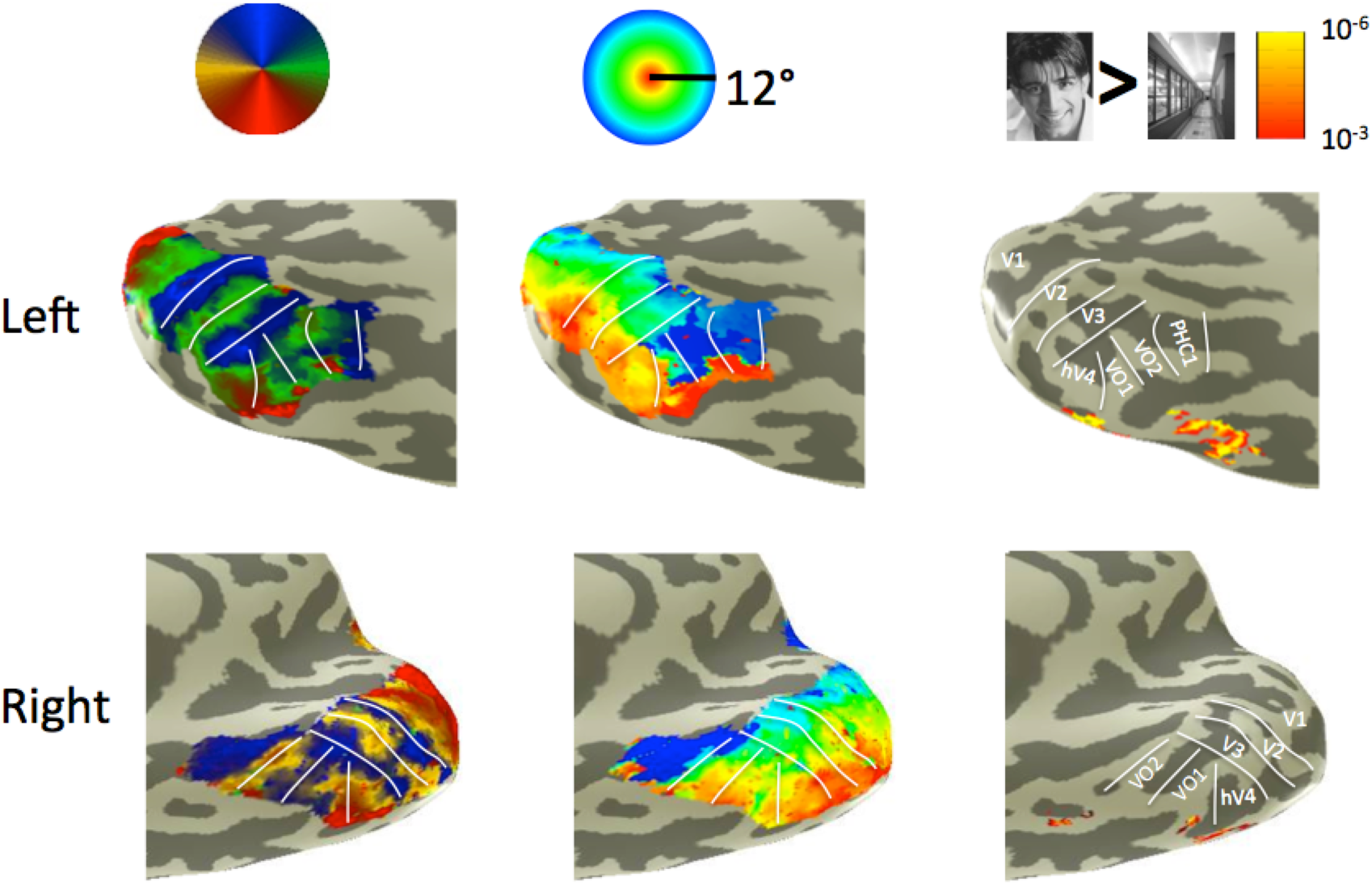
Developmental prosopagnosics show a typical functional organization of the ventral stream. *Left*: polar angle map; *Middle*: eccentricity map; *Right*: face-selectivity map. Maps display the ventral and medial inflated surfaces of posterior temporal and occipital cortices. Left and right hemispheres show data from two different DPs.

We compared the average volume of ROIs across groups, as well as the average variance explained by the pRF fit in each ROI. We did not find any significant differences in ROI volume or variance explained by the pRF model fit between groups (maximum difference in variance explained was 6% in hV4, *p* > 0.15; **Figure 2B**). The lack of significant differences in the volume, number, and anatomical arrangement of functional ROIs is consistent with previous findings showing that DPs have a normal functional organization of occipito-temporal visual cortex(15).

**Figure 2:**
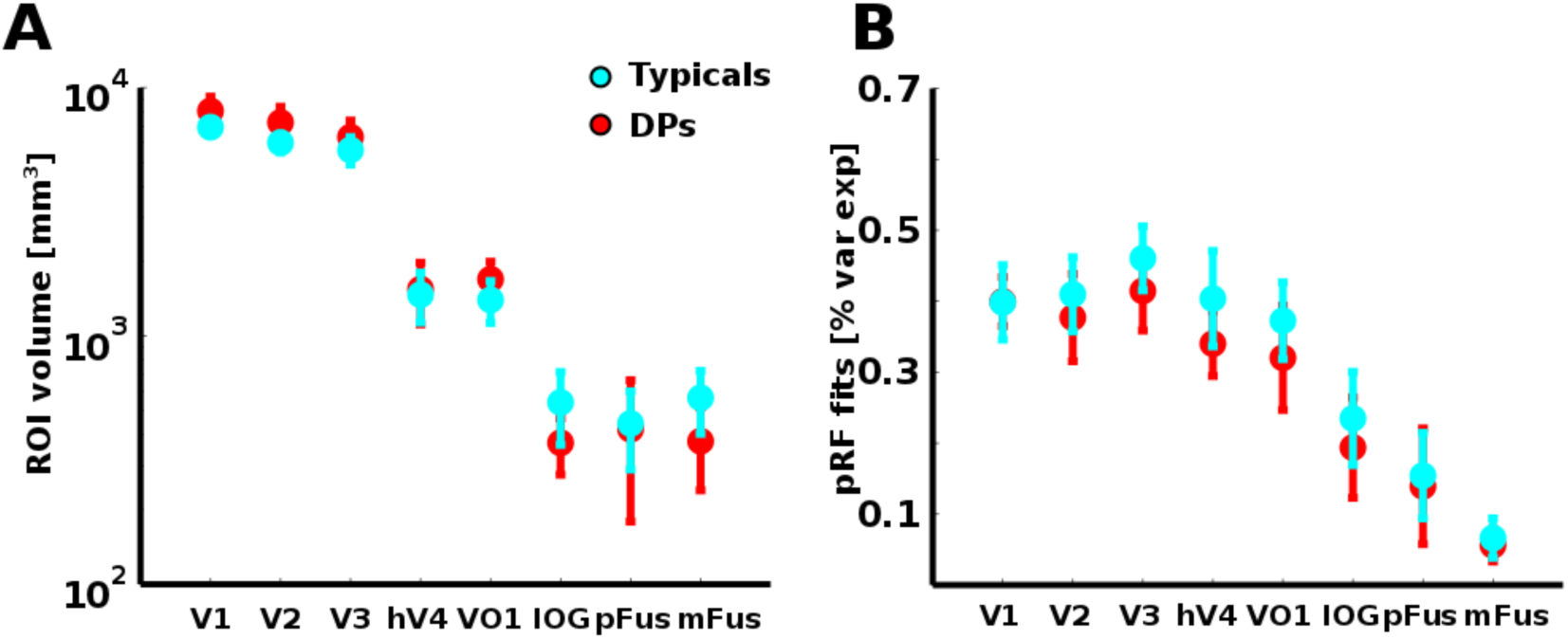
No significant differences of ROI sizes and pRF fits across DPs and typical participants. A. Average volume of visual field maps (V1-VO1) and face-selective ROIs (IOG, pFus, mFus). **B**. Average variance explained by pRF fits. Error bars are 95% confidence intervals (1.96 x standard error of the subject medians).

We tested for hypothesized differences in spatial processing between DPs and typicals by comparing the size and eccentricity of pRFs across the ventral visual cortex (**Figure 3A**). Where noted, data were combined across face-selective ROIs as this increases the reliability of the average of pRF parameters in each subject and accounts for some subjects’ lack of a particular ROI, as all have a combined face-selective ROI. We found no differences in pRF size between groups in V1 and V2. However, we found significant differences in the average pRF size in V3, hV4, and VO1, as well as in each of the face-selective regions (all *t*s>2.3, all *ps*<0.04). The average pRF size in hV4 and VO1 in typical adults was roughly double that of DPs (hV4: mean typicals = 2.5°±1.1°, mean DPs = 1.4°±0.42°; VO1: mean typicals = 4.4°±1.5°, mean DPs = 2.1°±1.4°), while pRFs in face-selective voxels were three times larger in typicals than DPs (mean typicals = 3.4°± 3.1°, mean DPs = 1°± 0.37°; combined face-selective ROI).

**Figure 3:**
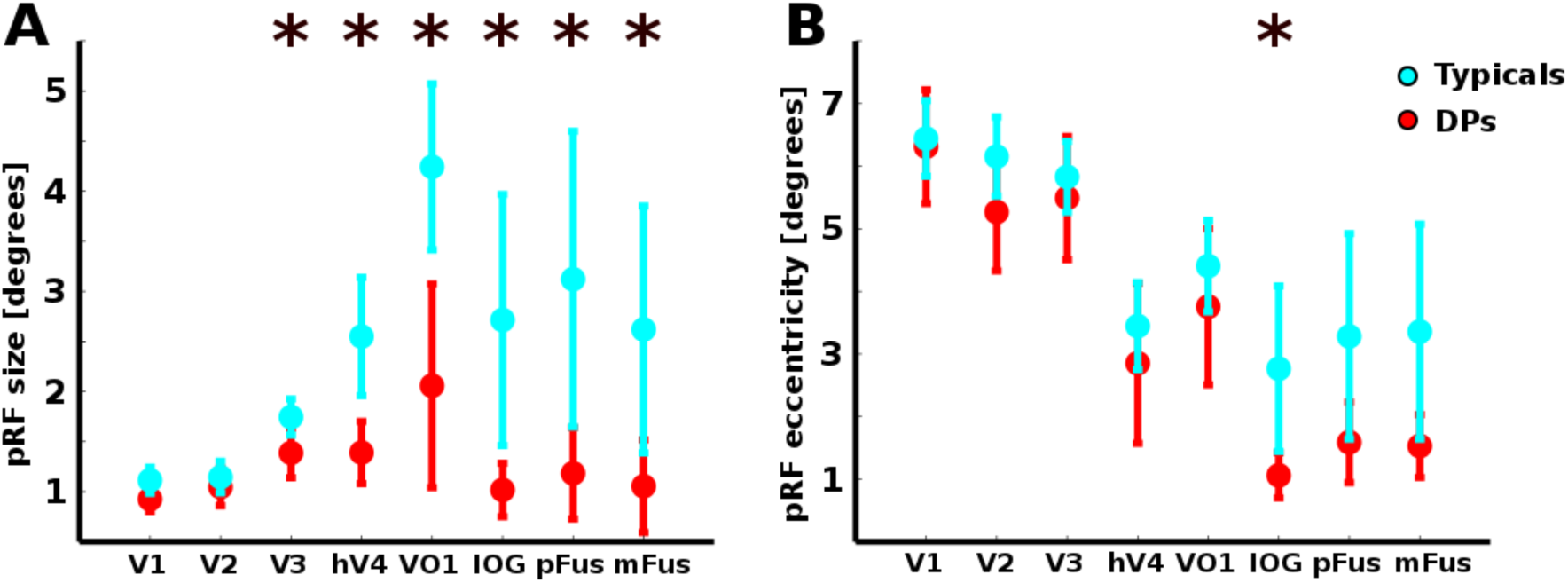
DPs have smaller pRFs in anterior visual field maps while face-selective areas have smaller pRFs and a tendency towards more foveal pRF centers. (A) DPs have smaller pRFs in all visual areas anterior to V2 **(B)** There is also a tendency for more foveal pRFs in face-selective regions. Asterisks represent p<0.05 from a two-tailed, unpaired t-test, without assumption of equal variance. Error bars are 95% confidence intervals (1.96 x standard error across subjects).

Consistent with previous findings, both groups show generally increasing pRF sizes moving from posterior to anterior visual field maps (e.g. V1 to VO1). Face-selective voxels in typical participants also show increased pRF size, with pRFs approximately three times larger than those in V1 (3.1° vs. 1.1°). However, in DPs, the average pRF size in face-selective regions was around 1°, showing almost no increase from pRF sizes in V1 (0.93°). We tested for a differential increase in pRF size across ROIs between groups using a repeated measures ANOVA with factors of group and ROI. We combined face-selective regions into a single ROI in each subject, as we did not observe differences across face-selective regions. We found significantly smaller pRFs in DPs relative to controls (main effect of group: F(1,20)=12.3, p=0.002) and expected differences in pRF size across ROIs (main effect of ROI, F(5,100)=18.3, p<10^−5^)). Notably, we found a significant group by ROI interaction (F(5,100)=3.9, p=0.026, sphericity corrected) showing that the difference in pRF size between the two groups systematically varied across ROIs. These results are consistent with our hypothesis that face processing deficits in DP are associated with smaller pRFs in face-selective regions. Moreover, reduced pRF size appears to be widespread across the anterior parts of ventral stream, affecting not only face-selective regions but also neighboring visual field maps hV4 and VO1.

A fundamental feature of visual cortex is that within an ROI, pRF size increases with eccentricity(30). Given the smaller pRFs observed in DPs, we examined if pRFs in DPs are closer to the fovea than in typicals. The average eccentricity of pRFs showed no significant differences between DPs and typicals in any of the visual field maps (max difference ~0.7° in V2, all *ps*>.15; **Figure 3B**). However, the average eccentricity of pRFs in DP’s face-selective regions ranged from 1.1° to 1.6°, less than half that of typicals (2.8-3.4°). PRFs were significantly more foveal in DPs in face-selective IOG (t(16)=2.44, p=0.03) and the combined face-selective ROI (t(17.9)=3.16, p=0.006), but marginal in pFus and mFus (ps<0.08). Comparison of pRF size as a function of eccentricity shows that in both groups, pRFs in each ROI increase in size from the fovea to the periphery (**Figure 4)**. Importantly, these plots also reveal that DPs have smaller pRFs in VO1 and face-selective regions across all eccentricities. This means that in face-selective regions, DPs have smaller pRFs than typicals at all eccentricities, as well as an increased foveal bias.

**Figure 4:**
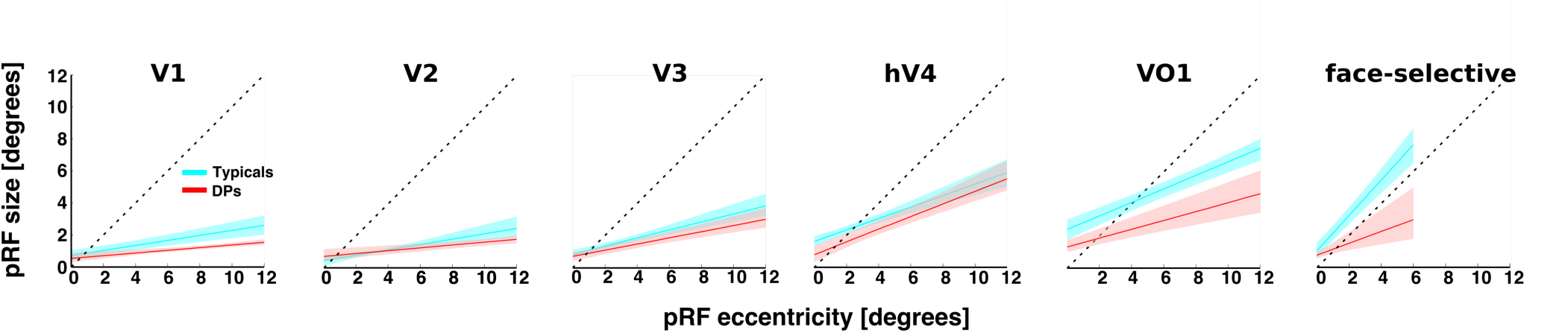
PRFs in DPs’ face-selective regions and VO1 are smaller at all eccentricities. Each panel shows pRF size as a function of the eccentricity of pRF centers for a given ROI. The dashed line corresponds to x=y, at which size scales exactly with eccentricity. Points above the dashed line have some ipsilateral visual field coverage. Lines are fit using a bootstrapping procedure across individual subject line fits. Bolded line is the median from the resulting distribution, and shaded area is the 95% confidence interval. Fits for the combined face-selective ROI were restricted to the central 6 degrees as there are few pRF centers outside that range (**Supp Fig. 4**). In DPs, like typicals, this function shows a linear relationship in each ROI, such that larger pRFs are more peripheral than foveal pRFs. However, DPs have smaller pRF size across eccentricities in VO1 and the combined face-selective ROI (95% confidence intervals never overlap).

*How do differences in pRF properties affect the visual field coverage across the two groups?* We combined the visual field coverage of all pRFs within an ROI to evaluate the coverage of the visual field obtained by the entire ROI. The visual field coverage of pRFs in early retinotopic areas V1, V2, and V3 showed the expected contralateral hemifield representation in both DPs and typicals (**Figure 5**). hV4 and VO1, show reduced visual coverage of the lower visual field in DPs compared to typicals, although DP’s visual field coverage is similarly contralateral, and extends to the periphery.

**Figure 5:**
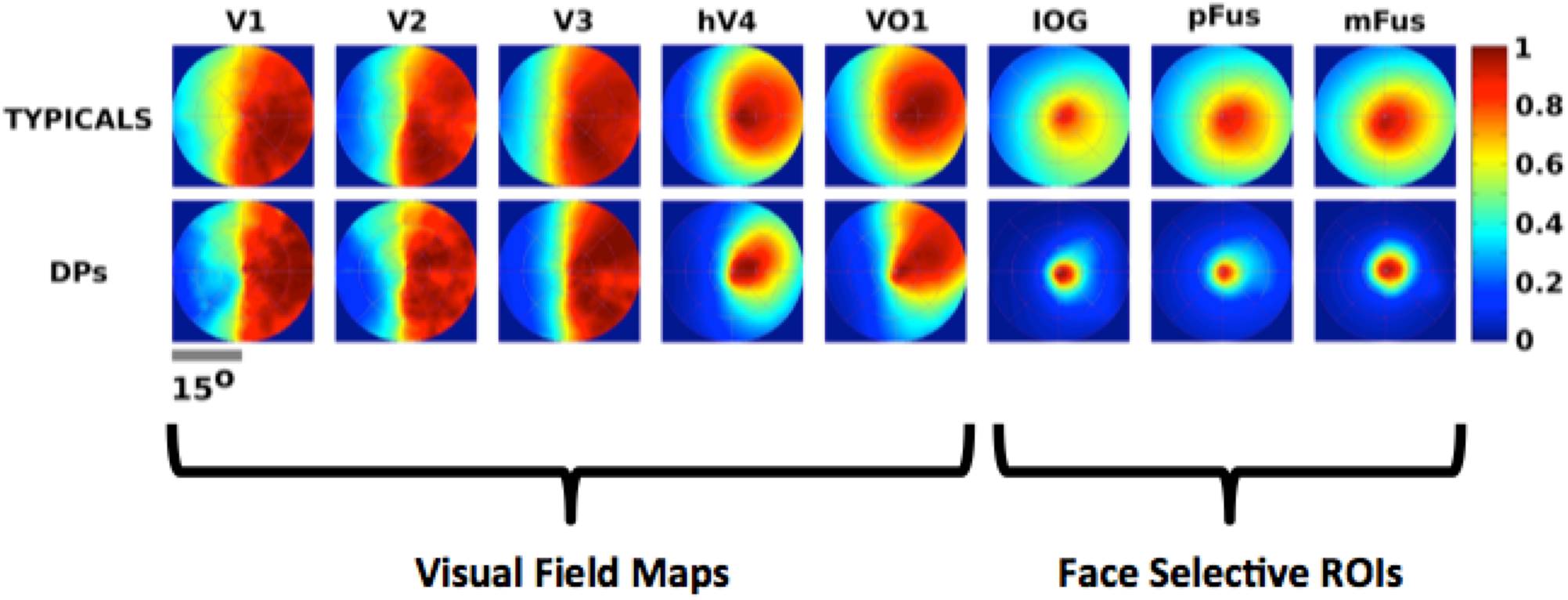
Visual field coverage of face-selective regions in DPs is restricted to the fovea. Panels display the across subject average maximum profile of pRF coverage of each retinotopic and face-selective region. Data from the right hemisphere are flipped across the vertical axis and combined with the left for each ROI. The color map represents a normalized group coverage of the visual field. In typicals (top row), the V1-V3 display the expected coverage of the contralateral hemifield, hV4 shows a bias towards the fovea, and VO1 showing a bias towards the upper visual field. Face-selective ROIs by contrast show a strong bias towards the fovea with some ipsilateral coverage. In DPs, V1-V3 coverage appears similar to typical adults, hV4 and VO1 show somewhat reduced coverage of the visual field, and the face-selective ROIs show substantially reduced coverage.

Visual field coverage in face-selective regions differs from the quarter or hemifield representations in visual field maps(33). In typicals, the visual field coverage of face-selective regions shows a bias towards the fovea and contralateral visual field, and but coverage also extends into the ipsilateral visual field (**Figure 5**). This ipsilateral coverage in face-selective regions is due to the fact that pRF sizes are larger than their eccentricity (**Figure 4**). In contrast, the combination of the smaller pRF sizes in face-selective regions and their concentration near the fovea in DPs affects their visual field coverage in a striking manner. Coverage does not extend to the ipsilateral visual field or the periphery, and is confined largely to the central 3° (**Figure 5**). These data show that face-selective regions of DP are especially devoted to the central portion of the visual field.

We hypothesized that variation in pRF size may be responsible for variation in face recognition ability, as smaller pRFs do not receive information from a large enough portion of the visual field to generate representations spanning multiple facial features. To test this hypothesis, we administered the Cambridge Face Memory Test (CFMT(36)) to all DPs (7/7) and available controls (10/15). The CFMT is widely used to identify DP subjects, and is correlated with measures of holistic face processing(37). We calculated correlations between pRF size, pRF eccentricity, and each participant’s performance on the CFMT.

Participant performance on the CFMT was significantly and positively correlated with pRF size in the combined face-selective ROI (r^2^=0.41, p<0.005; **Figure 6**) and VO1 (r^2^=0.40, p<0.007), but not in earlier visual areas (e.g., V1: r^2^=0.07, p>0.31). The correlation between CFMT score and pRF size was significantly larger in VO1 than V1 (one tailed t=−1.97, p<0.04) and marginally larger in the combined face-selective ROI than V1 (t=−1.69, p=0.06). As VO1 is not face-selective, the correlation between performance in the CMFT and pRF size in VO1 may at first appear surprising. However, pRF size is correlated between face-selective regions and VO1 (Supplemental Figure 3). pRF eccentricity did not predict CFMT performance in any ROI (maximum r^2^=0.04 in VO1, *ps*>0.44), although we did observe a trend for a positive relationship in the combined face-selective ROI (r^2^=0.19, p<0.08). In a model using both pRF size and eccentricity in the combined face-selective ROI to predict CFMT score, only pRF size significantly predicted behavior (size: t=2.6, p<0.03), and this combined model was not better than one using pRF size alone (F(14)=0.67, p>0.42).

**Figure 6:**
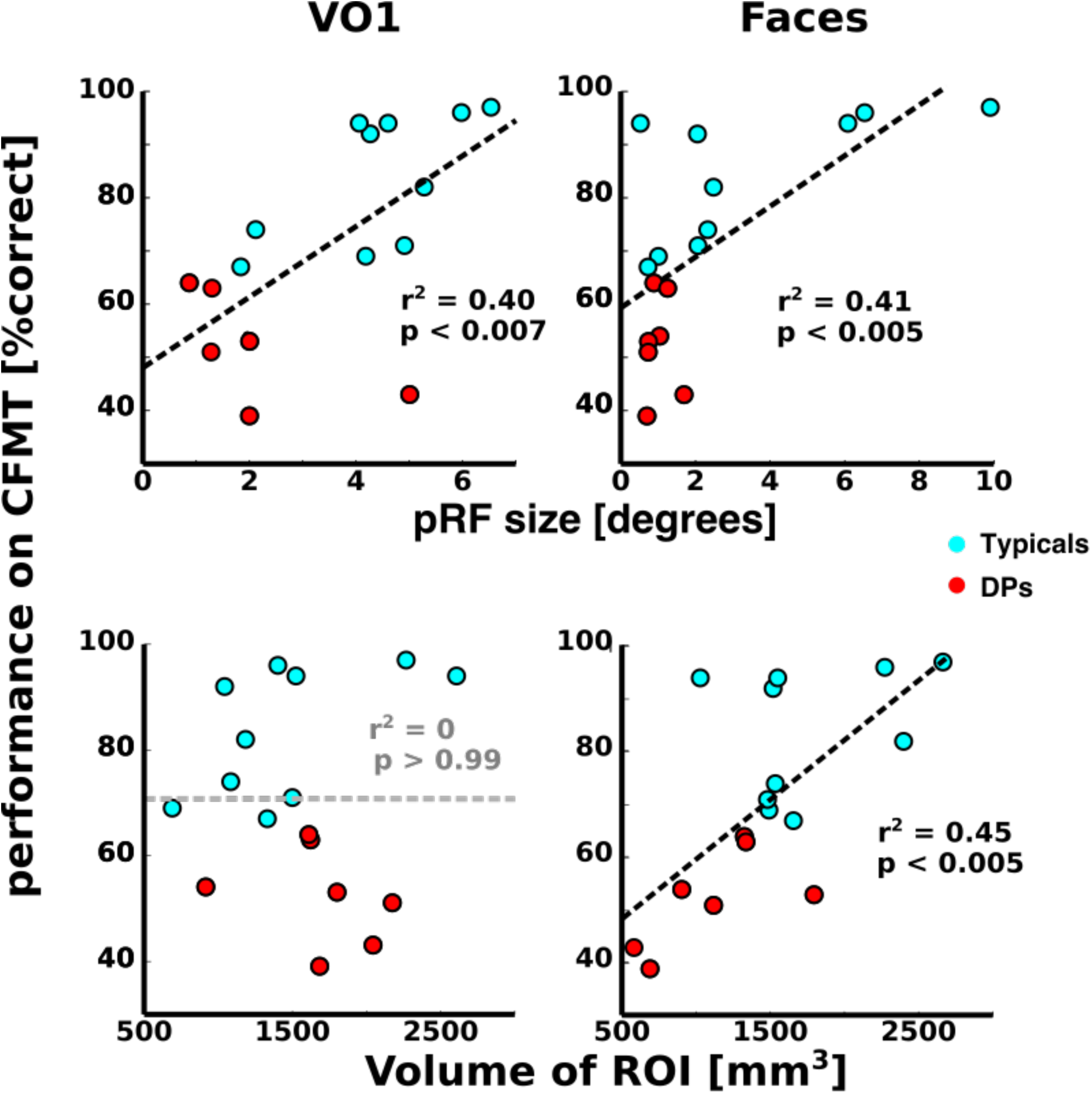
Relationships between performance in a face recognition task and ROI measures. *Top*: Correlation between accuracy on the Cambridge Face Memory Task (CFMT)(36) and the median pRF size in VO1, and the combined face-selective ROI. Each dot reflects a single subject. Significant correlations are shown in bold. *Bottom*: Correlation between the performance on the CFMT and the average ROI volume.

*How does our data fit with previous findings?* Prior work has shown that the volume of face-selective regions predicts performance on various face processing tasks in both DPs and typicals(14)(38)(39). We also found a positive relationship between performance in the CFMT and the volume of the combined face-selective ROI (r^2^=0.44, p<0.004; **Figure 6**). Otherwise, only V1 showed a negative correlation approaching significance (r^2^=0.22, p=0.055). Interestingly, in the combined face-selective ROI, pRF size and ROI volume are correlated (r^2^=0.47, p<0.005). Given the collinearity between pRF size and ROI volume, using both factors in a regression model does not significantly improve the model fit over using either alone. By contrast, VO1, shows a relationship between CFMT performance and pRF size, but shows no correlation between behavior and ROI size (r^2^=0, p>0.99), nor a correlation between ROI size and pRF size (r^2^=0.07, p>0.75).

## DISCUSSION

Measurements of pRFs across the ventral stream revealed that outside early visual cortex, DPs have smaller pRFs and restricted coverage of the visual field compared to typical adults. While we also found that DPs had more foveal pRF centers within the combined face-selective ROI, this did not explain the difference in size, as pRF sizes were smaller across eccentricities. Notably, our data show that accuracy in a face recognition task was positively correlated with pRF size in face-selective regions and VO1. These findings are consistent with the hypothesis that developmental prosopagnosia, at least in part, reflects a reduced access to spatially disparate information.

*What might be the source of reduced pRF size in DP?* The size of a voxel’s pRF reflects the combined output of many thousands of neurons, and it is impossible to disentangle the contribution of the size and scatter of neuronal receptive fields at the millimeter resolution afforded by fMRI. Nonetheless, the increase in pRF size as one moves from V1 to V2 to V3 and so on is thought to reflect the pooling of inputs from earlier visual areas(40)(41). Given the widespread differences in pRF size at later stages of the visual hierarchy between DPs and typicals, one possibility is that there is a generic difference between groups in the spatial pooling in the ventral processing stream.

Since DPs have smaller pRFs in much of ventral visual cortex, it is natural to ask if DPs exhibit behavioral problems beyond recognition of faces. In fact, some DPs have other subtle impairments, including discriminating objects(4)(5)(42)(43), navigation(44)(6)(45), perceiving biological motion(46), and holistic processing of non-face stimuli(47). While not all tasks have been tested in all DPs, differential impairment on non-face tasks is found both within(42)(46) and between studies(23)(28), suggesting the DP population is heterogeneous. This heterogeneity can be addressed with studies using larger sample sizes, and more complete behavioral characterizations.

The particular dependence of face processing on holistic representations(21) could also make it harder to develop compensatory strategies for face recognition, but allow for the development of useful alternative strategies for other kinds of visual tasks, such as object recognition or navigation. Affected individuals may therefore be less likely to report difficulties in other non-face domains, and differences in performance may be harder to assess experimentally. Future studies using larger sample sizes, and more complete behavioral characterizations can address the effect of smaller pRFs on processing of non-faces.

Our data also raise the possibility that DPs might recognize smaller faces more easily, since these would fit inside the covered part of the visual field at both the voxel and ROI level. We think this is unlikely. Spatial extent and position provide an important but limited characterization of the receptive field structure of face-selective voxels. If we think of the ‘true pRF’ of a face-selective voxel as a complete characterization of the relationship between possible stimuli and neural responses, it seems likely that other aspects of the pRF may be deficient in DP. Recent work showing that face-selective regions in DPs do not discriminate between intact and scrambled configurations of face parts is consistent with this idea(17).

While decreased pRF size in DPs is found outside face selective regions, there is a conjunction of neural differences in DPs that occurs only in face-selective regions. Only face-selective regions had reduced pRF sizes, increased foveal bias, restricted visual field coverage limited to the central visual field, and a reduction in total volume. It may be the combination of these factors and not just the reduction in pRF size generates the behavioral deficit in DP. In support of this possibility, we note that although the distributions of pRF sizes in ventral visual cortex strongly differed between DPs and typicals, some typical participants also had small pRFs (**Figure 6**). This suggests that small pRFs alone are not sufficient to produce deficits in face recognition.

One possibility is that the relationship between volume, pRF size, and CFMT performance has a developmental origin. Neuroimaging studies have reported the volume of face selectivity in ventral visual cortex expands from childhood, through adolescence, and into adulthood(38)(48)(49) and that this increase is associated with improved face recognition memory(38). These developmental data suggest the functional properties of brain regions supporting adult-like face recognition abilities change during normal development. One hypothesis is that only cortical regions with useful intrinsic properties will be recruited for face processing, such as regions with large receptive fields centered near the fovea(32). This proposal assumes that pRF properties in face-selective regions are relatively fixed across development. A second hypothesis is that pRF size and face selectivity may develop in tandem. Given that pRFs are also smaller in the adjacent visual field maps of DPs, this explanation is likely incomplete. Future research combining corresponding measurements of pRFs and face-selectivity in children can examine the validity of these new hypotheses.

In sum, we show that basic features of visual spatial processing as measured by pRFs may have behavioral consequences for high-level vision. These results are important for uncovering the neural basis of DP, and can serve as a stepping stone for studying other cognitive impairments which depend on integrating visual information across features and the visual field.

## METHODS

### Participants

Retinotopy and localizer data were obtained from 16 controls (8 Male; ages 18-39), and 9 DP s(3 male; ages 19-41). One DP fell asleep in three fMRI sessions. In another DP, we collected retinotopy data twice, but could not identify any visual field maps. This left 7 DPs for the analysis. We discarded data from one typical with noisy retinotopy and in whom we were unable to identify hV4 or VO1. DPs self-reported as having difficulty recognizing faces and scored more than two standard deviations below the population mean given for the Cambridge Face Memory Test(36) (Supplemental Figure 1). DPs also performed poorly on famous face recognition and recognition memory of unfamiliar faces, but normally in famous place recognition and recognition memory of unfamiliar places and objects (Supplemental Figures 1 and 2). Controls were all available subjects scanned on the same experiments on the same scanner with usable retinotopy. Some data from these participants has been previously published(34)(50). The available 10 of 15 control participants took the CFMT and scored in the typical range. All participants had normal or corrected-to-normal vision, were university students, or had at minimum an undergraduate degree, and provided written informed consent. Protocols were approved by the Stanford IRB on Human Participants Research.

### MR Data Acquisition

All MR data were collected at the Lucas Center for Imaging at Stanford University using a 3T GE Scanner. 2-4 T1-weighted SPGR scans (TR = 1,000 ms, flip angle = 45°, FOV = 200 mm, voxel size: 0.938 x 0.938 x 1.5 mm, 166 sagittal slices) were collected from each participant using an 8-channel whole brain coil. Anatomical images were averaged for each subject and segmented into white and gray matter using the Freesurfer auto-segmentation algorithm (http://freesurfer.net/) and hand editing in ITK-snap (http://www.itksnap.org/pmwiki/pmwiki.php). The gray-white boundary was used to create surface visualizations on inflated meshes using mrVista (https://github.com/vistalab/).

BOLD data were collected using 8-channel phased-array surface coil (Nova Medical, Inc. Wilmington, MA, USA). 32 slices oriented perpendicular to the calcarine sulcus were collected using a T2*-weighted gradient echo spiral pulse sequence with TR = 2,000 ms, TE = 30 ms, flip angle = 76°, FOV = 200 mm. Functional data was collected with 3.125 mm isotropic voxels with no gaps between slices. T1-weighted images with the same prescription as the functional scans were collected to facilitate alignment to each subject’s anatomy. Visual stimuli were projected onto a screen and viewed through an angled mirror mounted above the supine participant’s head.

### Retinotopy

Retinotopy stimuli were standard ring and wedge stimuli described previously(34). Subjects participated in two runs where we measured polar angle preferences using a 45° wedge rotating clockwise about fixation 22.5° every 2 s and two runs where we measured eccentricity responses employing a series of traveling concentric rings (1.4° in diameter) centered at fixation. The innermost ring consisted of a disk with a diameter of ~2.8° while the most peripheral extent of the checkerboard stimuli was ~14-15° from fixation. Both the wedge and ring stimuli contained 100% contrast black and white phase reversing checkerboards (at a rate of 4 Hz) and used six cycles (either a full rotation of the wedge or a full expansion of the ring) each lasting 32 s. Each run contained 6 cycles, which were interspersed with four 16-s blank periods (96 TRs in total). In addition, each scan began and ended with a blank 12-second block. Subjects were instructed to fixate on a central dot and press a button when the fixation color changed.

Population receptive fields (pRFs) were fit to the time series data of each voxel using the implementation in mrVISTA(30). For each voxel, the model estimates the pRF center in Cartesian coordinates and its size in degrees of visual angle. PRF fits in example voxels in left V2 and left IOG in a typical subject and a DP participant are shown in Supplementary Figure 9.

### Visual Field Map Definitions

V1-V3 were defined using widely agreed upon criteria. The dorsal and ventral arms of V2 and V3 were combined into a single ROI. hV4 and VO1 were drawn following the procedure outlined in Brewer et al 2005(35) and Witthoft et al(34). The foveal and peripheral boundaries of the visual field maps are limited by our stimuli in terms of their peripheral extent, and our ability to resolve polar angle in the fovea. Therefore, when we report ROI sizes for visual field map this reflects the extent of the visual field map between roughly 0-15°. Visual field delineations in all subjects are shown in Supplemental Figure 7.

### Category Localizer

All participants were presented with 7 classes of stimuli: adult male faces, child faces, indoor scenes, outdoor scenes, abstract objects, cars, and textures generated by scrambling the intact stimuli. All images were achromatic, square, and subtended approximately 11°. A run consisted of two presentations of each block type, except for the textures, which were seen four times per run. During each 12 s block, stimuli were shown at 1 Hz. Subjects were instructed to fixate and press a key when an image repeated. Image blocks alternated with 12 s fixation blocks. Each subject participated in two runs. Data were fit using a standard GLM analysis with boxcar regressors convolved with a hemodynamic response function. No smoothing was applied.

### Face-selective ROI definitions

Face-selective ROIs were defined in each individual using the contrast adult+child faces > indoor+outdoor scenes, with a voxel-wise threshold of p<0.001. This choice of contrast produces more widespread activations than other choices of contrast such as faces>objects(38), but our interest was in having as many voxels as possible for the pRF analysis. We defined three face-selective ROIs: one overlapping the inferior occipital gyrus just posterior and lateral to hV4 (IOG faces), and two on the fusiform (pFus-faces, and mFus-faces)(9). Face-selective ROIs in all subjects are shown in Supplemental Figure 7.

### ROI Analyses

A series of 3-way analyses of variance (ANOVAs, with factors of group x hemisphere x ROI) showed no effect of hemisphere on ROI size, pRF size, or variance explained by the pRF fit (all main effects and interactions with hemisphere as a factor *p*s>0.21). Therefore data were combined across hemispheres. Where mentioned in the text, data from the face-selective ROIs were aggregated to form a single face-selective ROI. This increased the robustness of our estimate of the median pRF size by having more voxels for each subject and ensuring that every subject had an ROI. ANOVAs comparing pRF size, variance explained, and eccentricity across the face-selective ROIs did not show significant differences within either group. It should be noted that the volume and variance explained for each ROI shown in figure 2 are un-thresholded.

For other analyses, all voxels with pRF fits explaining less than 5% of the variance were removed from the analysis. This value was chosen based on our estimate of a null distribution of the variance explained. In brief, we calculated the variance explained by the pRF fit in a region that has no retinotopic signal. Thus, we generated on each subject’s cortical surface a disk ROI of 5mm radius outside the visual field maps. This disk ROI was located in the parietal occipital sulcus of each subject, near where the POS and calcarine sulci meet and was roughly the size of pFus-faces. Results shown in Supplementary Figure 9 indicate that more than 95% of the voxels of this non retinotopic ROI are not well fit by the pRF model, as their variance explained is less than 5%. Therefore, we have chosen a liberal threshold for inclusion of voxels, which differentiates voxels that can be fit by a pRF model from those that cannot.

We also excluded voxels with pRF centers further than 15° from the fovea to eliminate pRFs with centers that are not on the display. The results are not dependent on the thresholding, as the same pattern of effects can be observed in the un-thresholded data, as well as data limited to pRFs that explain at least 10% or 20% of the variance (Supplemental Figures 4,5).

The across subjects average of variance explained, eccentricity, and pRF size for the pRFs in an ROI was calculated using the following procedure. First we obtained the median value of the parameter of interest across all voxels within that ROI for each subject, and then calculated the average and standard error of the participant medians for each group and ROI. All t-tests were performed on subject medians. Tests were two-tailed, unpaired, and did not assume equal variance. Similar results were obtained with the 10/15 typicals who took the CFMT (Supplemental Figure 6).

### Visual field coverage plots

Visual field coverage plots were generated for each subject using the following method. Each pRF was normalized to have a height of one. For all ROIs, the horizontal (x) position of pRF centers was flipped for the right hemisphere ROI and then, combined with pRFs from the left hemisphere. For example, all pRF centers in right V1 were flipped across the vertical meridian, moving them to the isomorphic location in the ipsilateral visual field. This produced a single bilateral ROI representation that maintains the lateralization of visual responses.

Coverage of the visual field by all pRFs for an ROI for an individual participant was defined as the maximum profile across the visual field of all the pRFs within that ROI. A 128x128 grid was created representing the visual field. For each voxel in the ROI we calculated the amplitude of the fit pRF at each location and took the maximum value found for each location in the visual field across all pRFs in the ROI. To reduce noise, 100 coverage plots were generated for each ROI for each participant using bootstrapping and averaged to create the participant’s visual field coverage for that ROI. Group average coverage plots were then created by bootstrapping across the subject averages. Similar results were obtained with the 10/15 typicals who took the CFMT (Supplemental Figure 7).

#### Size vs. eccentricity plots

The size by eccentricity plots for each ROI shown in **Figure 4** were generated using a bootstrap procedure. Data for each subject were separated into 0.5° eccentricity bins, and the average pRF size weighted by variance explained was calculated for each bin. On each of 10000 bootstraps, subjects from the appropriate group (either DP or typicals) were selected randomly with replacement and a line fit to the data. The fit line was evaluated at 0.1° intervals. The lines in Figure 5 are the 50^th^ percentile of the fit lines at each 0.1 degree interval, while the 95% confidence intervals are given by the 2.5^th^ and 97.5^th^ percentiles at each interval.

#### Correlation plots

For the correlation plots, the median value of pRF size and eccentricity for each subject within an ROI were used as distributions of these parameters can be non-normal. ROI volume is the total volume of the ROI without thresholding by variance explained by the pRF fits. These results were also robust across choice of threshold (Supplemental Tables 1-3).

